# KDM2B is a histone H3K79 demethylase and induces transcriptional repression via SIRT1-mediated chromatin silencing

**DOI:** 10.1101/228379

**Authors:** Joo-Young Kang, Ji-Young Kim, Kee-Beom Kim, Jin Woo Park, Hana Cho, Ja Young Hahm, Yun-Cheol Chae, Daehwan Kim, Hyun Kook, Sangmyung Rhee, Nam-Chul Ha, Sang Beom Seo

## Abstract

The methylation of histone H3 lysine 79 (H3K79) is an active chromatin marker and is prominant in actively transcribed regions of the genome. However, demethylase of H3K79 remains unknown despite intensive research. Here, we show that KDM2B (also known as FBXL10), a member of the Jumonji C family of proteins and known for its histone H3K36 demethylase activity, is a di- and tri-methyl H3K79 demethylase. We demonstrate that KDM2B induces transcriptional repression of *HOXA7* and *MEIS1* via occupancy of promoters and demethylation of H3K79. Furthermore, genome-wide analysis suggests that H3K79 methylation levels increase when KDM2B is depleted, indicating that KDM2B functions as an H3K79 demethylase *in vivo*. Finally, stable KDM2B-knockdown cell lines exhibit displacement of NAD^+^-dependent deacetylase SIRT1 from chromatin, with concomitant increases in H3K79 methylation and H4K16 acetylation. Our findings identify KDM2B as an H3K79 demethylase and link its function to transcriptional repression via SIRT1-mediated chromatin silencing.

## Introduction

Chromatin structure is modulated by diverse covalent histone modifications (Cheung et al., 2000; Strahl and Allis, 2000) Combinations of such modifications direct both global and specific transcriptional outcomes (Berger, 2007; Lee et al., 2010). Among these modifications, histone lysine methylation is linked to both activation and repression of transcription. As in the case of most epigenetic modifications, histone methylation and demethylation are dynamically regulated by histone methyltransferases (HMTases) and demethylases (Martin and Zhang, 2005; Mosammaparast and Shi, 2010; Wysocka et al., 2005).

Histone H3 lysine 79 methylation (H3K79me) is catalyzed by the HMTase disruptor of telomeric silencing-1 (Dot1)-Like (DOT1L), which is the mammalian homolog of the yeast Dot1 (Feng et al., 2002; Lacoste et al., 2002; Singer et al., 1998; van Leeuwen et al., 2002). DOT1L is considered the only H3K79 HMTase in mammals. However, recent reports suggest that the nuclear SET (Su(var)3-9, Enhancer-of-zeste, Trithorax) domain (NSD) family of HMTases, including response element II binding protein (RE-IIBP) (also known as NSD2), has H3K79-methylating activities (Morishita et al., 2014; Woo Park et al., 2015). On the basis of analyses of its crystal structure, H3K79 is a surface-exposed residue and is located in close proximity to lysine 123 of histone H2B in yeast cells (Luger et al., 1997; Ng et al., 2002b). Regulation occurs by trans-crosstalk, and ubiquitylation of histone H2BK123 (H2BK123ub) is necessary for H3K79me formation (Briggs et al., 2002).

H3K79 methylation is linked to active gene transcription (Ng et al., 2003; Okada et al., 2005; Okada et al., 2006; Schubeler et al., 2004; Vakoc et al., 2006). In addition, H2BK123ub, H3K4me, H3K79me, and RNA polymerase II (Pol II) phosphorylation occurs sequentially during transcriptional elongation (Nguyen and Zhang, 2011). H3K79me plays a role in DNA repair through interaction with Rad9/53BP1, and the levels of H3K79me change during the cell cycle (Feng et al., 2002; Huyen et al., 2004; Schulze et al., 2009; Wysocki et al., 2005). During embryogenesis, H3K79me regulates the expression of developmental genes (Ooga et al., 2008; Shanower et al., 2005). In addition, aberrant hypermethylation of H3K79 results in the activation of oncogenes and leukemic transformation (Bernt et al., 2011; Bitoun et al., 2007; Mueller et al., 2007).

The Jumonji C (JmjC) domain-containing histone demethylase, KDM2B, also known as FBXL10, preferentially demethylates both H3K36me2/3 and H3K4me3 (Frescas et al., 2007; Tsukada et al., 2006; Wang et al., 2011). KDM2B also mediates the monoubiquitylation of the histone H2AK119 as a component of noncanonical polycomb-repressive complex 1 (PRC1) in embryonic stem cells (ESCs) (Farcas et al., 2012; Wu et al., 2013). KDM2B enhances reprogramming of ES cells via its binding to unmethylated CpG sites through the zinc finger (ZF)-CXXC motif. CpG recognition and PRC1 targeting by KDM2B are important for the deposition of H2AK119ub1 and for further recruitment of PRC2 to a subset of CpG islands, which is an activity limited to variant PRC1 complexes (Blackledge et al., 2014; He et al., 2013; Liang et al., 2012).

KDM2B contributes to the development of tumors *in vivo*, probably via H3K36 demethylase activity. Overexpression of KDM2B inhibits cellular senescence by repressing the mouse loci p16Ink4a, p19Arf, and p15Ink4b, and by repressing the human loci retinoblastoma (Rb) and p53, resulting in cellular immortalization (Pfau et al., 2008; Tzatsos et al., 2009). In addition, wild-type KDM2B but not a mutant with defective demethylase activity enhances the progression of pancreatic cancer in a mouse model (Tzatsos et al., 2013). Furthermore, H3K36 demethylase activity is required for leukemic transformation in a Hoxa9/Meis1–induced mouse bone marrow transplantation (BMT) model (He et al., 2011).

In this study, we identified KDM2B as a histone demethylase that can catalyze the removal of di-and tri-methyl groups from the H3K79 lysine residue. We also found that KDM2B induces SIRT1- mediated chromatin silencing by removing H3K79me, which leads to transcriptional repression.

## Results

### Identification of the H3K79me2 peptide-interacting proteins

In an attempt to identify a potential H3K79 demethylase, we hypothesized an interaction between H3K79me site and a corresponding demethylase or a certain complex with H3K79 demethylase activity. This hypothesis led us to perform a peptide pull-down assay using H3K79me2 peptides with K562 nuclear extracts. By liquid chromatography-tandem mass spectrometry (LC-MS/MS), we showed that methylated H3K79 was associated with chromobox homolog 8 (CBX8), one of the various types of CBX proteins directing the canonical PRC1 complex as an interacting component (Figure 1A and B). To rule out the possibility of non-specific binding, we carried out pull-down assays between CBX8 and H3K79me0 peptides as negative controls, and confirmed that CBX8 interacted with H3K79me2 peptides but not with H3K79me0 peptides (Figure 1C). PRC1, one of the polycomb group (PcG) multiprotein complexes, has a diverse composition that depends on the presence or absence of CBX proteins (Di Croce and Helin, 2013; Gil and O’Loghlen, 2014). Previous studies isolated nuclear proteins bound to KDM2B and identified the BCL6 co-repressor (BCOR) complex by mass spectrometry (Sanchez et al., 2007). Another studies identified the presence of the KDM2B-containing non-canonical PRC1-BCOR-CBX8 complex (Beguelin et al., 2016; Sanchez et al., 2007). To investigate the associations of these proteins, we performed immunoprecipitation (IP) experiments and showed that KDM2B bound CBX8 (Figure 1D). On the basis of these results, we suggest that CBX8 is associated with KDM2B in a certain complex and that KDM2B possibly functions at the H3K79me site.

**Figure 1.**
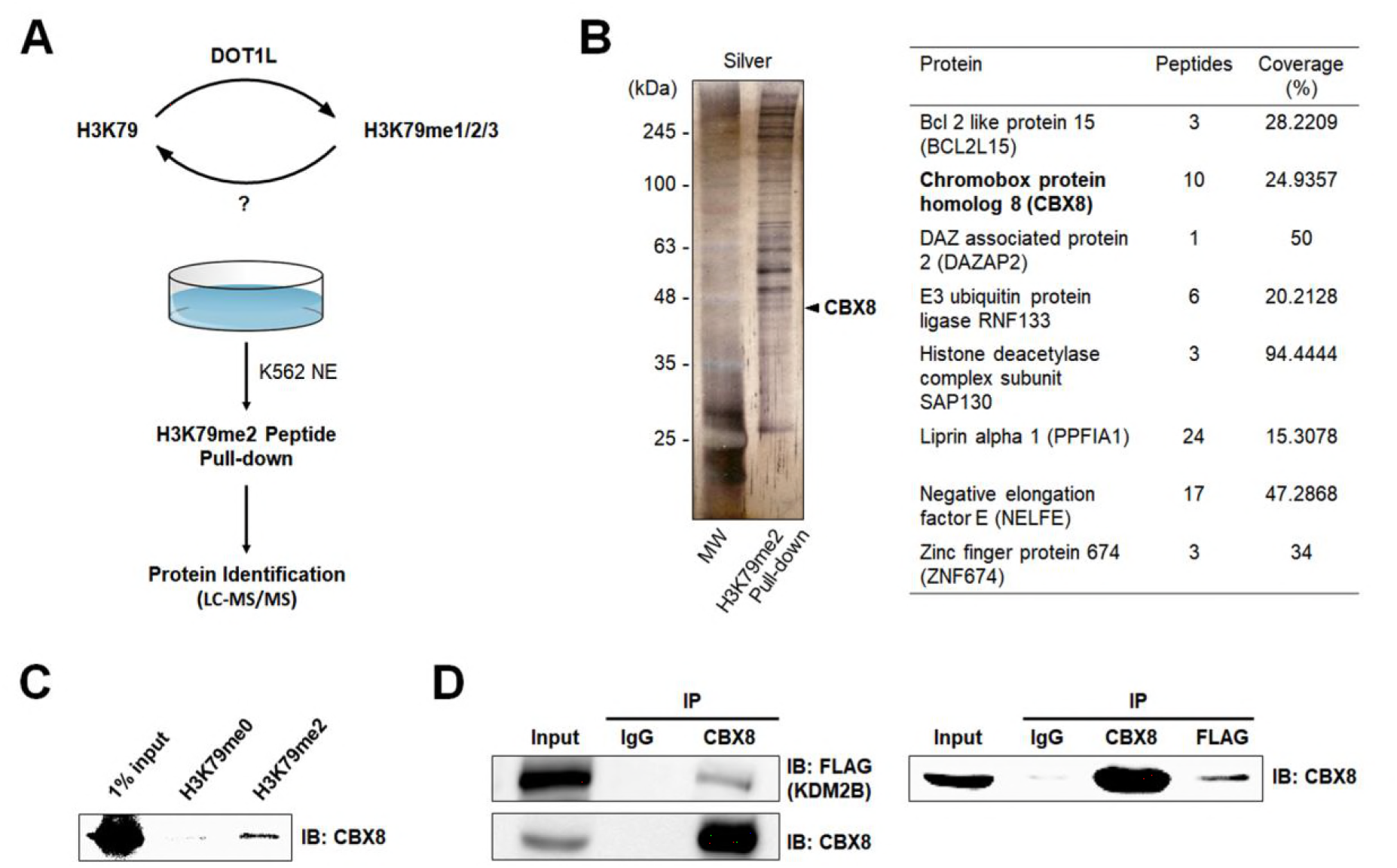
Identification of H3K79me2-associated proteins. **(A)** Schematic representation of pull-down assay using H3K79me2 peptides in K562 nuclear extract. **(B)** The pulldown complex was separated using 12% SDS-PAGE and visualized by silver staining. The gel was sliced into three sections and subjected to LC-MS/MS analysis. Several peptide sequences containing CBX8 fragments were detected. **(C)** Pull-down of H3K79me2 and H3K79me0 (as a negative control) in K562 whole cell lysate. **(D)** Immunoprecipitation of endogenous CBX8 and ectopically overexpressed FLAG-KDM2B in 293T cells.

### KDM2B demethylates H3 di-/tri-methyl-K79 in *vitro* and *in vivo*

In addition to being a subunit of the non-canonical PRC1 complex, KDM2B is the only known demethylase involved in other types of complexes containing CBX8, including the BCOR complexes. Therefore, we decided to test the H3K79 demethylation activity of KDM2B (Figure 2-figure supplement 1A). Ectopic expression of FLAG-tagged full-length KDM2B in 293T cells markedly reduced the levels of H3K79me3 and, to a lesser degree, reduced the levels of H3K79me2. No decreases in H3K79me1 levels were observed (Figure 2A, left panel). As expected, KDM2B also decreased the levels of H3K36me2. However, a mutant form with a substitution of histidine with alanine in the JmjC domain catalytically deficient in H3K36 demethylase activity, KDM2B-H242A, failed to demethylate H3K79me3, H3K79me2, and H3K79me1 (Figure 2A, left panel). We observed that DOT1L rescued the effects of KDM2B on H3K79 but not on H3K36, which confirmed the H3K79 demethylase activity of KDM2B (Figure 2-figure supplement 1B).

**Figure 2.**
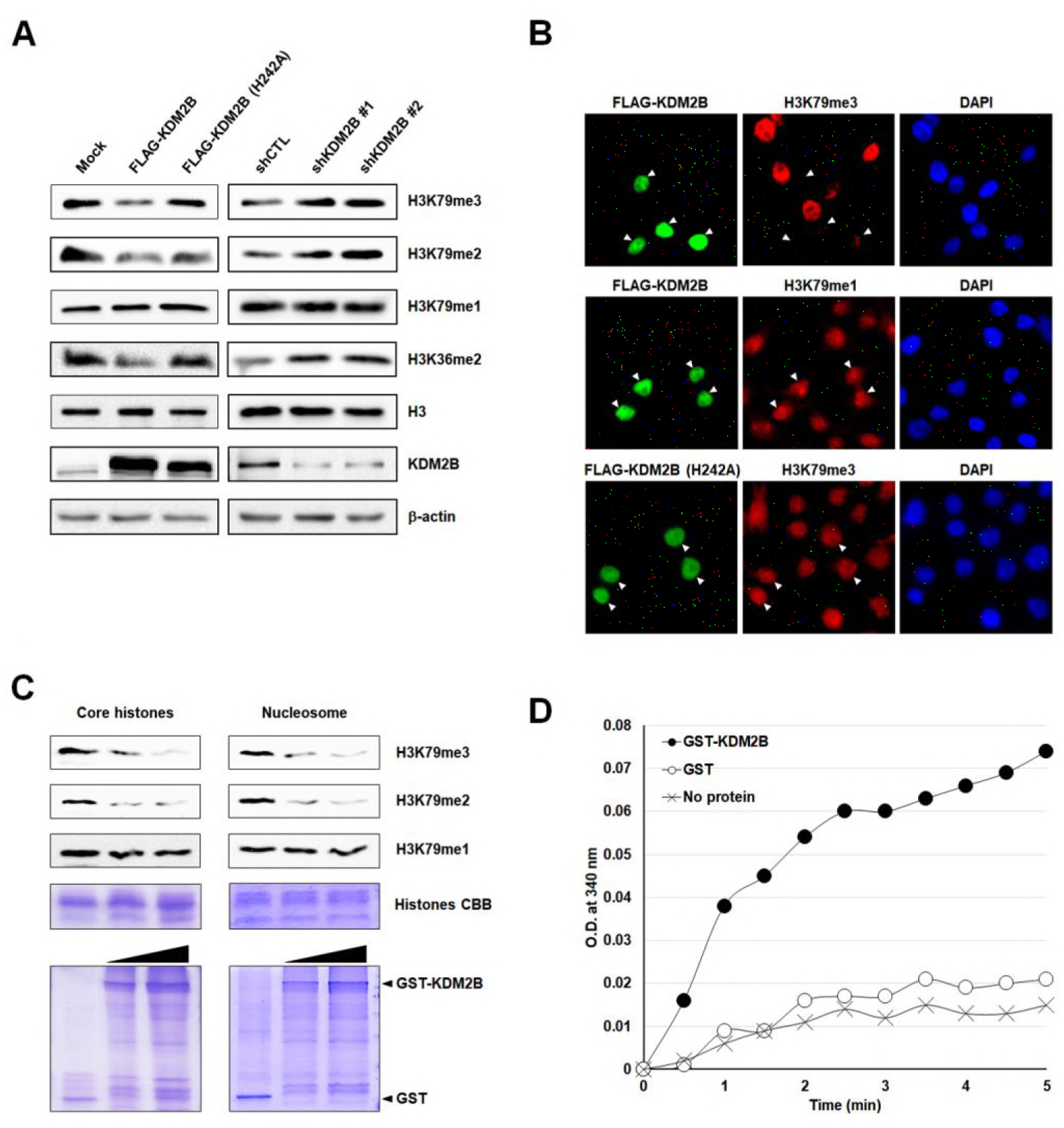
KDM2B demethylates histone H3K79me2/3. **(A)** Effects of overexpression and knockdown of KDM2B on H3K79 demethylation. FLAG-tagged KDM2B wild-type and catalytically deficient H242A mutant were transfected into 293T cells. H3K79me1/2/3 levels were detected by western blotting. KDM2B knockdown stable 293T cells were analyzed in the same way. **(B)** Immunocytochemistry of 293T cells expressing FLAG-KDM2B wild-type and KDM2B-H242A. Changes in signal intensity were detected after double immunostaining with anti-FLAG and anti-H3K79me1/3 antibodies. Arrows indicate transfected cells. **(C)** KDM2B *in vitro* demethylase assay. Upper: Core histones and nucleosomes were used as substrates. Demethylation activity of KDM2B against di- and tri-methyl H3K79 is shown. Lower: Purified GST and GST-KDM2B_1–734_ recombinant proteins used were visualized by Coomassie staining. **(D)** Formaldehyde production was detected by FDH assay. NADH production was measured at OD 340 nm after 3-hour reaction using GST-KDM2B together with recombinant MLA histone H3K79me3 as substrates. Absence of demethylase and GST were used as negative controls.

We next investigated the H3K79 demethylation at the global level by comparing methylation status in stable KDM2B knockdown 293T cells. We used two independent shRNAs targeting different regions of KDM2B: one in the coding sequence (CDS) and the other in the 3’-untranslated region (UTR). We found that RNA interference (RNAi) of endogenous KDM2B led to increased H3K79me3 and H3K79me2 levels, but mono-methylation was not affected (Figure 2A, right panel). Furthermore, immunocytochemistry showed that overexpression of KDM2B in 293T cells resulted in a loss of H3K79me3, in contrast with the strong H3K79me3 staining signals observed in adjacent non-transfected cells (Figure 2B). However, overexpression of KDM2B had no detectable effect on H3K79me1 levels that were determined by immunocytochemistry (Figure 2B).

We confirmed the H3K79 demethylase activity of KDM2B through *in vitro* assays. It was necessary to decide whether the above observations derived from indirect effects of genetic manipulation or from the direct enzymatic activity of KDM2B. After incubation of core histones with increasing concentrations of GST-KDM2B containing the JmjC, CXXC, and plant homeodomain (PHD) domains (amino acids 1–734), we observed reduced levels of H3K79me2 and of H3K79me3 (Figure 2C, left panel). We examined the effects of KDM2B on nucleosomes. Similar to the effects on core histones, KDM2B demethylated both H3K79me2 and H3K79me3 of nucleosomes (Figure 2C, right panel). We concluded that KDM2B had H3K79 demethylase activity both *in vitro* and *in vivo*. We performed isothermal titration calorimetry (ITC) experiments to test binding affinity of KDM2B towards tri-methylated H3K79 peptides. Given that H3K79me3 had a much lower dissociation constant (Kd) than H3K79me0, GST-KDM2B_1–734_ bound to H3K79me3 with stronger affinity (Figure 2-figure supplement 2A). To demonstrate the specificity toward H3K79, we assessed whether KDM2B removes the methyl groups from recombinant histone H3 with methyl-lysine analogs (MLAs) which was specifically methylated by chemical alkylation reaction (Jia et al., 2009; Simon et al., 2007), using formaldehyde dehydrogenase (FDH) assays. FDH assay measures the production of formaldehyde, a by-product derived from demethylation reaction, by monitoring the reduction of nicotinamide adenine dinucleotide (NAD+) into NADH (Lizcano et al., 2000; Shi et al., 2004). We used fluorescence-based detection method and assessed demethylase activity of KDM2B. It was observed that KDM2B effectively produced marked amounts of formaldehyde from MLA histone H3 containing tri-methylation on K_c_79, confirming that H3K79 is a substrate for demethylation catalyzed by KDM2B (Figure 2D). The FDH assay system also worked for MLA histone H3K_c_36me2 as a substrate (Figure 2-figure supplement 2B). Furthermore, we performed *in vitro* histone demethylase assays and LC-MS/MS analysis with MLA histone H3K_c_79me3. Incubation with GST-KDM2B_1–734_ led to demethylation of MLA histone H3K_c_79me3 and to increase of H3K_c_79me1 level (Figure 2-figure supplement 2C). Western blot analysis of MLA histone H3K_c_79me3 incubated with GST-KDM2B_1–734_ confirmed a marked decrease in H3K79me3 level (data not shown). These results indicate that KDM2B is an H3K79me2/3-specific demethylase.

### Structural analysis of H3K79 demethylation by KDM2B

To obtain insights into the mechanism of the demethylation reaction, we modeled H3K79 onto the KDM2B structure, based on the H3K36-bound KDM2A structure that was previously determined (Cheng et al., 2014). Given that the location of H3K79 within the protein structure is in the loop between the two alpha-helices, this region could be accessible to the active site of KDM2B with a slight conformational change. Remarkably, the sequences around the K79 and K36 of histone H3 are similar in their chemical properties. In accordance with the sequence alignment of residues surrounding the H3K79 and H3K36 (Figure 3A), this modeling exercise indicated that the hydrophobic residues in the H3K79 peptide (Phe78, Leu82, and Phe84) are involved in hydrophobic interactions with KDM2B. This is similar to what occurs in the interactions of the H3K36 peptide with the complex structure of KDM2A (Figure 3B). The residues (His242 and His314) for binding of iron and alpha-ketoglutarate (α-KG) were also conserved in the modeled structure (Figure 3B), indicating concomitant bindings of these co-factors with the H3K79 peptide. Together, these findings indicate that H3K79 is a plausible demethylation site for KDM2B.

**Figure 3.**
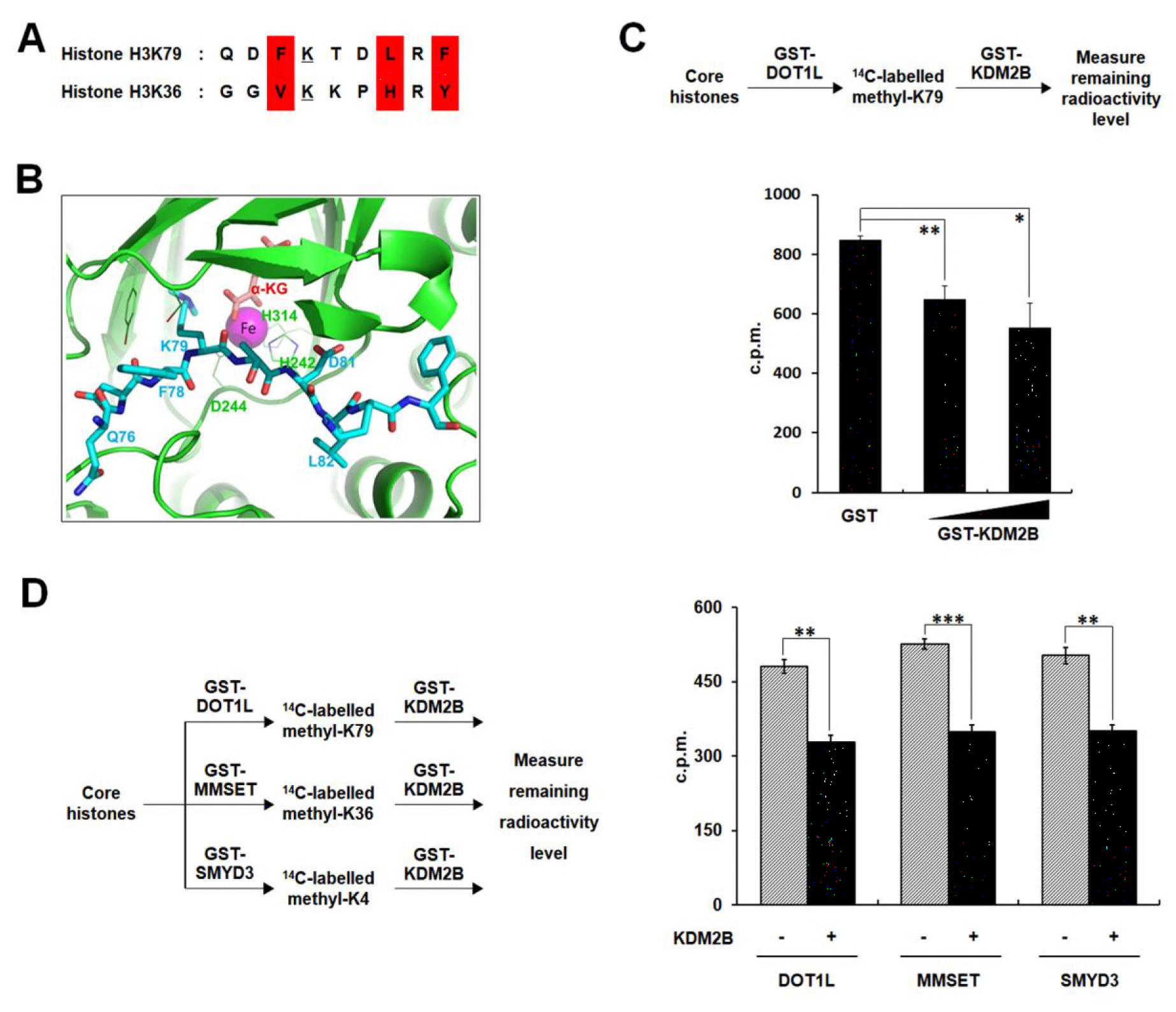
Structural modeling of KDM2B demethylase activity. **(A)** Comparative sequence analysis of nine amino acids around K36 and K79 of histone H3. **(B)** Crystal structure of KDM2B with enlarged residues critical for binding to H3K79. Iron co-factor and H3K79 residues in the catalytic pocket are shown. **(C)** Radioactive assay using core histones. H3K79 residues were ^14^C-labeled by GST-DOT1L through histone methyltransferase assay and the histones were used in histone demethylase assays with GST-KDM2B. Remaining radioactivity levels from histones were measured by scintillation counting to assess demethylase activity. All error bars indicate standard error of mean (SEM) for at least triplicated experiments. **(D)** Radioactive assays using core histones. H3K79, H3K36, and H3K4 residues were ^14^C-labeled by DOT1L, MMSET, and SMYD3, respectively. ^14^C-labeled histones were used in histone demethylase assays with KDM2B. Remaining radioactivity levels from histones were measured by scintillation counting to assess demethylase activity. All error bars indicate SEM for at least triplicated experiments.

We also verified the H3K79 demethylase activity of KDM2B, using core histone substrates radiolabeled by DOT1L (which is responsible for methylation at K79 sites). KDM2B significantly decreased the levels of H3K79-methylated histones (Figure 3C). In addition, since KDM2B was well known as H3K36 and H3K4 demethylase (Frescas et al., 2007; Tsukada et al., 2006; Wang et al., 2011), we compared the activities of KDM2B between H3K79, H3K36, and H3K4, using ^14^C-labeled histones by DOT1L, multiple myeloma set domain (MMSET), and SET and MYND domain containing protein 3 (SMYD3), respectively (Figure 3-figure supplement 1). We detected considerable reductions in radioactivity of each histone substrate following demethylation by KDM2B (Figure 3D). Therefore, these findings suggest that H3K79 demethylase activity of KDM2B is catalytically effective based on the degrees of H3K36 and H3K4 demethylation.

### KDM2B induces transcriptional repression

Since H3K79me was involved in transcriptional activation (Ng et al., 2003; Okada et al., 2005; Okada et al., 2006; Schubeler et al., 2004; Vakoc et al., 2006), we speculated that demethylation of the site by KDM2B might induce transcriptional repression. To elucidate the effects of KDM2B on target gene expression, we first tested whether KDM2B specifically downregulated transcription of the two best-studied mixed lineage leukemia (MLL) fusion target loci, the *HOXA7* and *MEIS1*, preferentially regulated by changes in H3K79me level. Luciferase reporter assays showed that KDM2B overexpression resulted in *HOXA7* and *MEIS1* transcriptional downregulation, and, conversely, KDM2B knockdown resulted in their upregulation (Figure 4A). These results are consistent with the previously examined repressive function of KDM2B. To show whether the enzymatic activity of KDM2B was required for the repressive function of the protein, the catalytically inactive mutant (H242A) and ΔCXXC mutant were tested in reporter assays using the *HOXA7* promoter. A significant absence of transcriptional repression was observed in assays using the ΔCXXC mutant, and partial repression was observed in assays using the H242A point mutant (Figure 4B). To examine whether the demethylation of H3K79 catalyzed by KDM2B led to downregulation of general transcription, we carried out luciferase assays using simian virus 40 (SV40) and thymidine kinase (TK) promoters. Transient overexpression of KDM2B repressed luciferase activity on both promoters, whereas KDM2B depletion resulted in transcriptional activation (Figure 4-figure supplement 1A). We next investigated whether transcriptional activities of KDM2B and DOT1L were in opposite direction through the regulation of H3K79me. Knockdown of DOT1L recapitulated KDM2B-mediated transcriptional repression on the same two promoters (data not shown).

**Figure 4.**
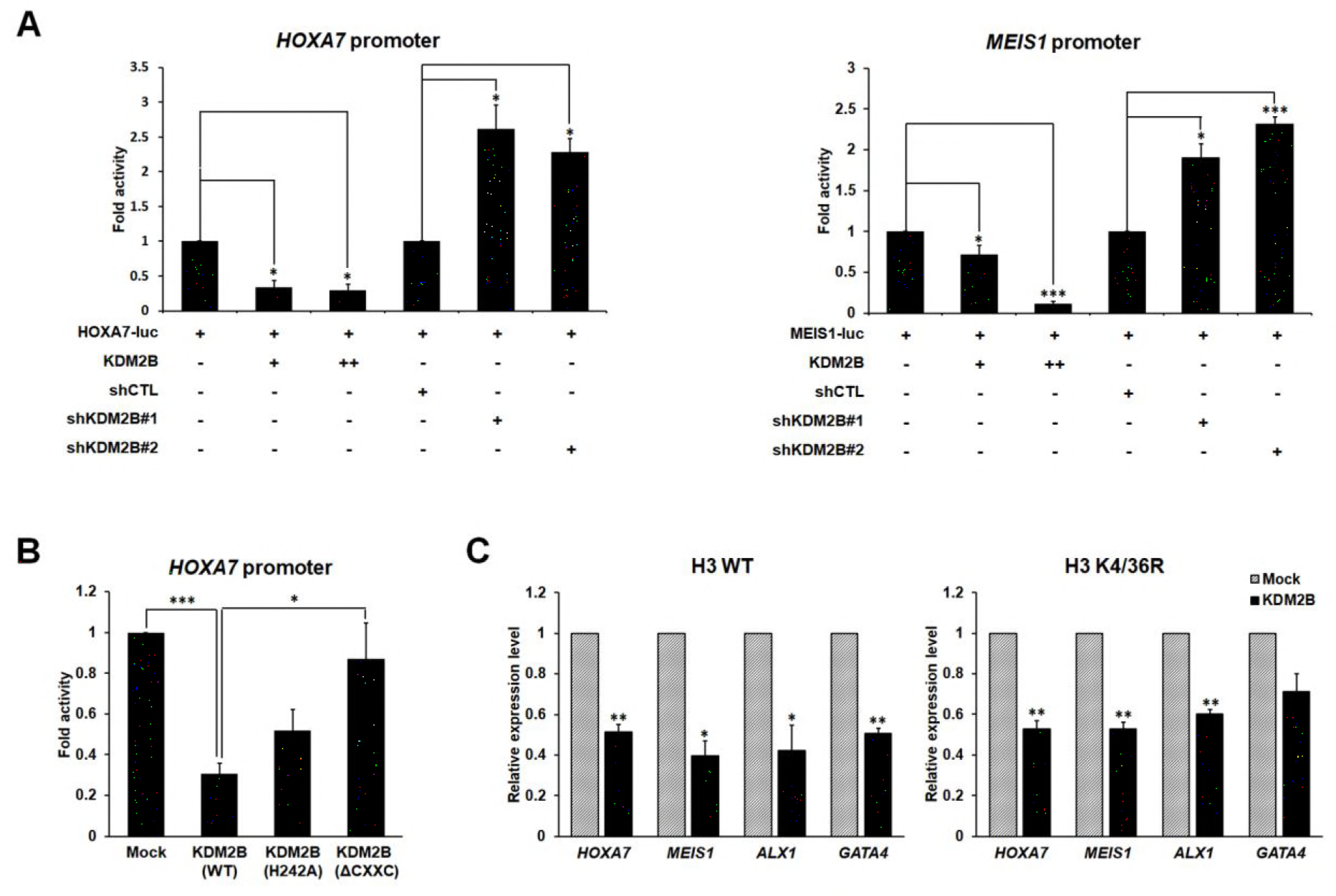
KDM2B-mediated transcriptional repression via H3K79 demethylation. **(A)** Luciferase assay using KDM2B target gene promoters. Promoter activities were analyzed with overexpression or depletion of KDM2B. All values are shown as standard error of mean (SEM), n = 3; \**p* < 0.05, \*\**p* < 0.01, \*\*\**p* < 0.001. **(B)** Luciferase assays using catalytically inactive mutants and deletion mutants of KDM2B. All values are shown as SEM. **(C)** qRT-PCR analysis of target gene expression in cells stably overexpressing histone H3 wild-type or K4/36R mutant. All error bars indicate SEM for at least triplicated experiments.

Based on published data, we selected four genes from approximately 2,500 genes directly targeted by KDM2B. *HOXA7*, *MEIS1*, *ALX1*, and *GATA4* have been reported as KDM2B and PcG target genes by chromatin immunoprecipitation (ChIP)-sequencing (seq) and ChIP analyses using mouse ESCs (Blackledge et al., 2014; Bracken et al., 2006; Farcas et al., 2012; Wu et al., 2013). Using these four known target genes, we demonstrated that H3K79 demethylation alone, without H3K36 or H3K4 demethylation, was involved in KDM2B-mediated repression. We measured the transcription levels of the four target genes using wild-type and K4/36R mutant histone H3. As expected, these four genes showed clear downregulation in H3 K4/36R-overexpressing stable cells, similar to their levels in H3 wild-type-overexpressing stable cells when KDM2B was ectopically expressed (Figure 4C). Taken together, we established that H3K79 demethylation is indeed required for KDM2B-mediated repression in addition to H3K36 and H3K4 demethylations.

### KDM2B depletion regulates gene expression through genome-wide accumulation of H3K79me

To determine the global location of H3K79me sites under the influence of KDM2B, we performed ChIP-seq in KDM2B knockdown stable 293T cells. A comparison of the two ChIP-seq data sets (control shRNA (shCTL) vs. shKDM2B) using heat maps showed that, at the global human genome level, H3K79me3 was enriched under the condition of KDM2B depletion (Figure 5A). The normalized sequencing depth of ChIP-seq data revealed that, under the KDM2B knockdown condition, methylation marks were considerably enriched around the centers of H3K79me3 peaks, suggesting that KDM2B ablation strongly upregulated H3K79me3 (Figure 5B). We confirmed that marked increases in H3K79me3 occupancy (at the genome-wide level) were correlated with KDM2B knockdown by analyzing the mean ChIP-seq tag density of shKDM2B compared to that of a shCTL (Figure 5C). ChIP-seq binding profiles on individual genes such as *GATA4* and *PDE3B* clearly indicated that H3K79me3 levels were upregulated in the shKDM2B stable cell line (Figure 5D). To validate the ChIP-seq experiment, we selected a KDM2B-responsive gene, *PDE3B*, and performed ChIP-qPCR. H3K79me3 accumulated on the *PDE3B* promoter when KDM2B was depleted (Figure 5E). To characterize chromatin profiles near transcription start sites (TSSs) of H3K79me3-occupied genes, sensitive to KDM2B depletion, we compared H3K79me3 peaks with published ChIP-seq studies for KDM2B in human acute myeloid leukemia (AML) cells (van den Boom et al., 2016). We noticed that alterations of H3K79me were correlated with KDM2B binding. KDM2B peaks were localized at *EGLN1* and *CYFIP1* loci, and H3K79me3 occupancy increased at these two genes after KDM2B depletion (Figure 5-figure supplement 1A). 61% of genes having more than 2-fold change in H3K79me3 levels overlapped with KDM2B target genes (Figure 5-figure supplement 1B). The genes whose H3K79me3 levels increased under the shKDM2B condition were selected and categorized by gene ontology (GO) term analysis (Figure 5-figure supplement 2). Notably, these genes were related to well-known functions of H3K79me, including transcriptional regulation and cell cycle control. These results suggest that KDM2B is responsible for a considerable amount of the H3K79 demethylation that occurs on its target genes.

**Figure 5.**
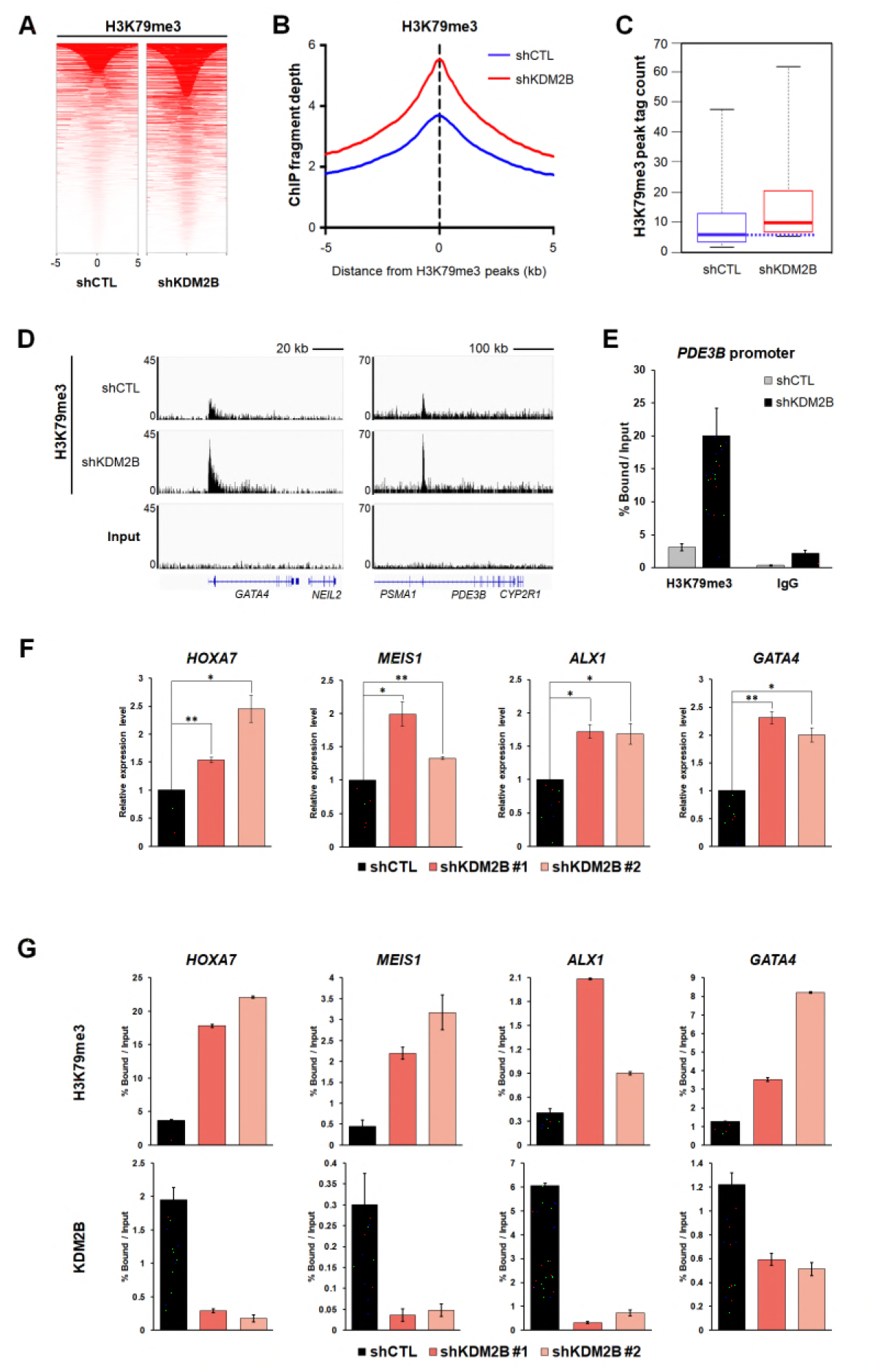
KDM2B lowers H3K79me3 occupancy at target gene promoters. **(A)** ChIP-seq heat map covering 10 kb regions across H3K79me3 peaks. **(B)** H3K79me3 ChIP-seq signal normalized by input sample in the control (blue line) and shKDM2B cells (red line). **(C)** Change in H3K79me3 occupancy following KDM2B knockdown. **(D)** Input and H3K79me3 ChIP-seq profiles over two regions of the genome in control and shKDM2B cells. Below the sequencing traces, *GATA4* and *PDE3B* genes are indicated. **(E)** H3K79me3 mark at *PDE3B* promoter in KDM2B knockdown cells analyzed by ChIP-qPCR. **(F)** qRT-PCR analysis for detection of target gene expression. KDM2B was depleted in 293T cells. **(G)** ChIP-qPCR analysis for measurement of KDM2B enrichment and H3K79me3 levels on target gene promoters. KDM2B was stably knocked down in 293T cells. Promoters of *HOXA7*, *MEIS1*, *ALX1*, and *GATA4* were immunoprecipitated and amplified using specific primers.

We next sought to understand the effects of KDM2B recruitment on the levels of H3K79me3 on target gene promoters. We performed ChIP and quantitative real-time PCR (qRT-PCR) analyses. Overexpression of wild-type KDM2B considerably decreased H3K79me3 levels on the promoters of four genes tested and led to transcriptional repression (Figure 5-figure supplement 3A and B). By contrast, the catalytically deficient KDM2B mutant failed to change H3K79me levels and gene expression, although it was still recruited to the target promoters (Figure 5-figure supplement 3A and B). Using two independent KDM2B-knockdown stable cell lines, we observed clear increases in H3K79me3 deposition on the *HOXA7*, *MEIS1*, *ALX1*, and *GATA4* promoters and concomitant induction of these target genes expression (Figure 5F and G). We obtained similar results when the ChIP experiments were normalized to histone H3 occupancy (Figure 5-figure supplement 4A). Also, the normalized ChIP results for H3K36me2 and H3K4me3 as well as H3K79me3 support the working model of KDM2B-mediated demethylation in a way of targeting multiple sites (Figure 5-figure supplement 4B). Collectively, these data show that KDM2B is recruited to specific target gene promoters and demethylates H3K79me3, resulting in downregulation of transcription.

### Recruitment of SIRT1 protein to target gene promoters by KDM2B

Previous studies showed that H3K79me by Dot1 regulated gene silencing (Ng et al., 2002a; Singer et al., 1998; van Leeuwen et al., 2002). Silent information regulator (SIR) proteins preferentially bind chromatin that contains hypomethylated H3K79, and they block H3K79me (Ng et al., 2003). As a member of the sirtuin family of proteins, SIRT1 is the human homolog of the yeast Sir2 protein, and mediates deacetylation of histones H3, H4, and H1 (Vaquero et al., 2004). To delineate the relationship between H3K79 demethylation and SIR protein-mediated transcriptional repression, we tested whether KDM2B induced the recruitment of SIRT1 to the target gene promoters. We measured the levels of SIRT1 and H4K16ac on the *HOXA7*, *MEIS1*, *ALX1*, and *GATA4* promoters by ChIP-qPCR analysis in KDM2B knockdown stable cell lines. We found that depletion of KDM2B increased H3K79me3 levels (Figure 5G), leading to disruption of chromatin binding of SIRT1 and to increase in H4K16ac levels (Figure 6A). Enrichment of H4K16ac under KDM2B ablation provides evidence for a correlation between H3K79me and H4K16ac in transcriptional upregulation. A recent study on the functional interplay between DOT1L and bromodomain-containing protein 4 (BRD4) reported that H3K79me, by itself, was not necessary for transcriptional activation, and that H4 acetylation was required for the actual regulatory effects to occur as downstream events (Gilan et al., 2016). To evaluate a model of SIRT1 recruitment, we overexpressed KDM2B in the KDM2B knockdown stable cell line. As expected, H3K79 hypomethylation-facilitated SIRT1 binding was rescued by ectopic KDM2B (Figure 6-figure supplement 1). We tested whether KDM2B interacted with SIRT1. IP analysis in 293T cells overexpressing KDM2B and SIRT1 showed that KDM2B bound SIRT1, and the interaction between endogenous SIRT1 and KDM2B was also detected (Figures 6B and C). SIRT1 has been shown to inhibit chromatin binding of DOT1L by H3K9me2 accumulation and chromatin compaction via SUV39H1 localization (Chen et al., 2015). To examine whether the decreases in H3K79me levels mediated by KDM2B were due to its association with SIRT1 and concomitant failure of DOT1L recruitment, we performed IP assay in wild-type and catalytic mutant of KDM2B overexpressed cells, and confirmed that KDM2B-H242A also bound SIRT1 (Figure 6-figure supplement 2). Given that KDM2B-H242A did not alter H3K79me3 levels, as shown in Figure 2A and Figure 5-figure supplement 3B, we concluded that H3K79me reduction by KDM2B did not originate from DOT1L inaccessibility caused by KDM2B-SIRT1 interaction. This interaction suggests that KDM2B induces SIRT1 recruitment possibly via H3K79 demethylation and facilitates H4K16 deacetylation. To demonstrate the role of SIRT1 recruitment in transcriptional repression by KDM2B, we tested whether the SIRT1 inhibitor (sirtinol) abolished KDM2B-mediated repression. Treatment with sirtinol resulted in a marked recovery of transcription of KDM2B target genes, despite of KDM2B overexpression (Figure 6D). This is consistent with the results of a previous study that described the requirement for H4 acetylation in transcriptional activity via alteration of H3K79me levels (Gilan et al., 2016). Taken together, these results indicate that H3K79 demethylation by KDM2B and subsequent tethering of SIRT1 are both involved in KDM2B-mediated transcriptional repression.

**Figure 6.**
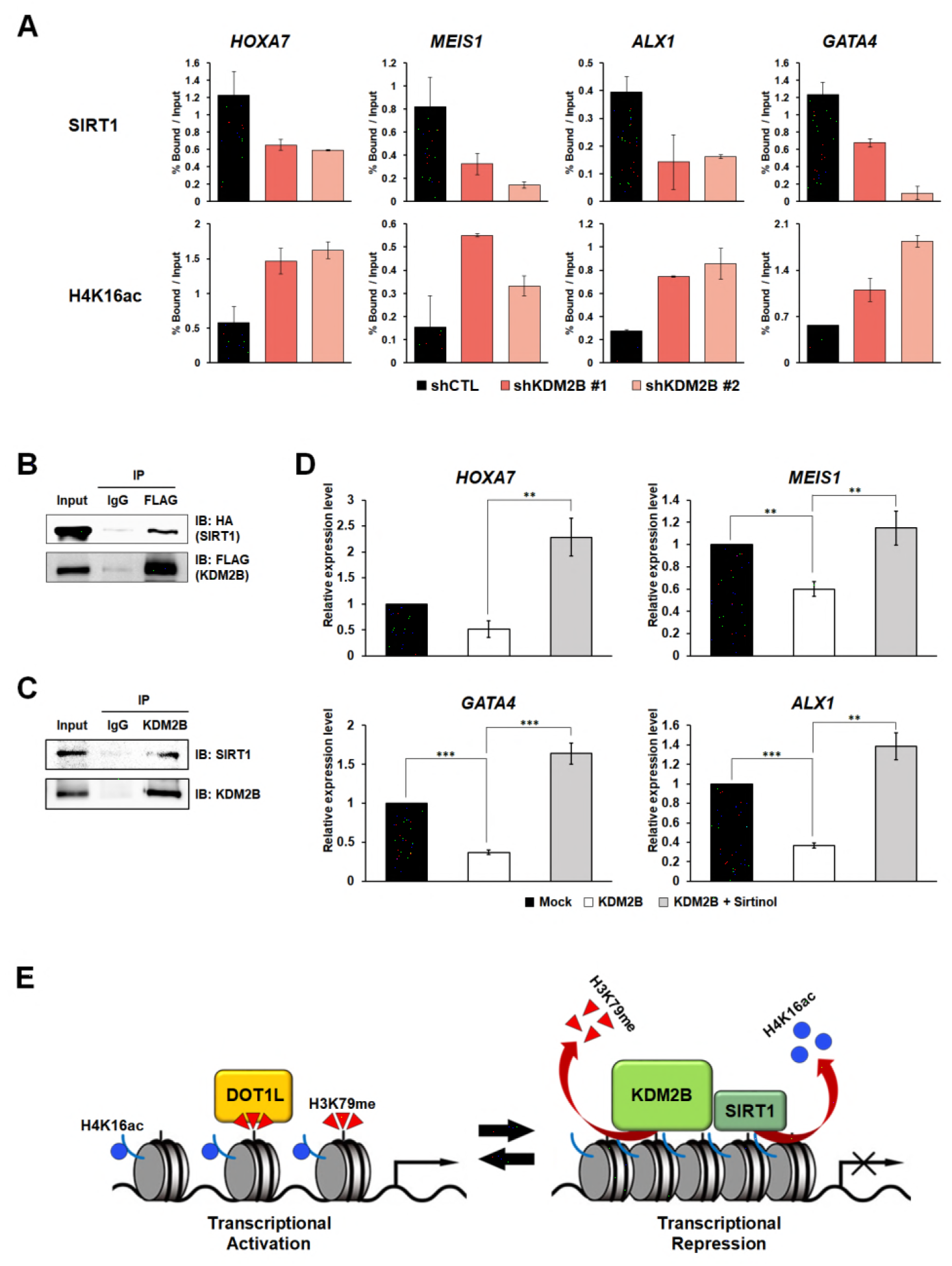
KDM2B-mediated H3K79 demethylation induces recruitment of SIRT1 to chromatin. **(A)** ChIP analysis was performed using anti-SIRT1 and anti-H4K16ac antibodies in stable KDM2B knockdown 293T cells. The immunoprecipitated DNA fragments were analyzed by qRT-PCR from the promoter regions of *HOXA7*, *MEIS1*, *ALX1*, and *GATA4*. **(B)** IP of ectopically overexpressed FLAG- KDM2B in 293T cells transfected with FLAG-KDM2B and HA-SIRT1. **(C)** IP of endogenous KDM2B in 293T cells. **(D)** qRT-PCR analysis for detection of target gene expression. 293T cells were treated with DMSO or sirtinol for 2 h following KDM2B overexpression. All error bars indicate SEM for at least triplicated experiments. **(E)** Proposed model of transcriptional repression of target genes by KDM2B via H3K79 demethylation and SIRT1 recruitment.

## Discussion

Although H3K79 is located in the globular domain of histone H3, it is localized on the surface of the nucleosomal structure, and can be accessed by epigenetic modifiers (van Leeuwen et al., 2002). The fact that H3K79me is evolutionarily conserved in a wide variety of eukaryotes is a strong indication of its fundamental role in the regulation of chromatin structure (Mersfelder and Parthun, 2006). Although extensive studies have been conducted to characterize the role of H3K79me in transcriptional regulation and its physiological outcome, the identity of the H3K79 demethylase has been uncertain. In this work, we identified KDM2B as a histone H3K79me2/3 demethylase. By pull-down assay followed by mass spectrometry analyses, we identified H3K79me2-interacting proteins, including CBX8. As a component of canonical PRC1 complex, CBX8 recognizes the H3K27me3 site and interacts with RING1B to contribute to H2AK119ub and maintenance of transcriptionally repressive state (Di Croce and Helin, 2013; Gil and O’Loghlen, 2014). The non-canonical PRC1-BCOR-CBX8 complex represses bivalent promoters and contains KDM2B as a component (Beguelin et al., 2016). In contrast, CBX8 interacts with MLL-AF9 and TIP60, playing a role in transcriptional activation of MLL-AF9 target genes (Tan et al., 2011). It is possible that CBX8 regulates gene expression in both positive and negative ways, depending on the enzymatic properties of binding partners, by serving as a molecular adaptor that localizes catalytic subunits in the complex to target genes. The reason for the interaction between H3K79me2 and CBX8 protein requires further investigation.

KDM2B is a member of the F-box protein family that includes KDM2A, which also shows H3K36 demethylase activity. KDM2B and KDM2A share conserved domains, including the JmjC and CXXC domains, and both proteins bind CpG islands (Blackledge et al., 2010). They both catalyze the demethylation of H3K36me2, which is a highly abundant modification and is specifically depleted at CpG islands (Blackledge et al., 2010). According to studies of different H3K27 demethylase activities of UTX and UTY (Agger et al., 2007; Hong et al., 2007), it remains to be investigated whether KDM2A also catalyzes H3K79 demethylation. KDM2A does not associate with PcG proteins, and its function in transcriptional regulation differs subtly from that of KDM2B.

H3K79me is found throughout the euchromatic regions of the genomes of yeast and higher eukaryotes, but it is significantly under-represented in silent chromatin (Ng et al., 2003). Sir2, which is involved in heterochromatin formation, is not recruited to the region of the chromosome containing H3K79me in yeast (Ng et al., 2002a). H3K79me2 is excluded from telomeres and from mating-type loci, to which Sir2 is recruited and mediates heterochromatin formation (Im et al., 2003; van Leeuwen et al., 2002). H3K79me2/3 are also associated with active gene expression in yeast and mammalian cells (Im et al., 2003; Pokholok et al., 2005). A recent study suggested that inactivation of DOT1L in MLL-AF9 leukemia cells enhanced SIRT1 occupancy at most genes marked with H3K79me, demonstrating an antagonistic relationship between DOT1L and SIRT1 (Chen et al., 2015). In fact, the heterochromatin proteins Sir3 and Dot1 compete for the basic patch of histone H4 in yeast. H4K16ac by Sas2 displaces Sir3 and enables Dot1 to bind H3K79, methylating H3K79 (Altaf et al., 2007; Millar et al., 2004).

Our current study reveals that KDM2B is an H3K79me2/3 demethylase and acts as a transcriptional co-repressor. By ChIP-seq and ChIP-qPCR analyses, we determined that KDM2B led to loss of H3K79me on target genes. Interestingly, previous study found that knockout of DOT1L reduced H3K79me2 globally, but it resulted in only a subset of H3K79me2/3 marked loci to be downregulated (Bernt et al., 2011). In addition, the effects of individual epigenetic modifiers on gene expression were much more specific and limited than predicted by changes in the status of a single histone modification (Lenstra et al., 2011). On the basis of these studies, we concluded that even though KDM2B influenced H3K79me at a genome-wide level, additional gene-dependent and context-dependent mechanisms might be involved in the regulation of gene expression. Recent studies demonstrated the role of histone H4 acetylation as a link between H3K79me and transcriptional activation (Gilan et al., 2016). This suggested the importance of a particular chromatin context in the regulatory processes mediated by changes in H3K79 methylation. We provided evidence that KDM2B induced the recruitment of SIRT1 to target gene promoters, leading to H4K16 deacetylation and chromatin silencing. This is consistent with an earlier report that CBX8 is a binding partner of SIRT1, and that they cooperatively mediate transcriptional repression (Lee et al., 2013).

To rule out the hypothesis that KDM2B-mediated repression in our study was derived from demethylation of two previously known substrates of KDM2B, H3K4 and H3K36, we overexpressed K4/36R double mutant histone H3 that is not subjected to methylation or demethylation. Transient overexpression of KDM2B following selection of H3 K4/36R-overexpressing stable cells enabled us to rule out repressive effects of K4 and K36 demethylations as the explanation and enabled us to propose an independent mechanism via a new target residue, K79. The wide histone substrate repertoire for KDM2B demethylase activity, which includes H3K36, H3K4, and H3K79, is interesting and suggests the possibility that KDM2B may also have non-histone protein substrates. In addition, considering the identification of new H3K79 HMTases other than DOT1L, it is reasonable to speculate that there may be other H3K79 demethylases besides KDM2B yet to be identified.

Given that the H3K36 demethylase activity of KDM2B was not necessary for its variant PRC1- mediated H2AK119ub1 and transcriptional repression, we proceeded to determine whether H2AK119ub1 was required to enable the H3K79 demethylase activity of KDM2B in the variant PRC1 complex. To test this, we used PRT4165, a potent inhibitor of PRC1-mediated histone H2A ubiquitylation. H2AK119ub levels decreased in the presence of PRT4165, but H3K79me3 demethylation catalyzed by KDM2B was not altered, suggesting that the H3K79 demethylase activity of KDM2B was independent of PRC1-mediated H2AK119ub1 (data not shown). Via the demethylation of H3K79, however, KDM2B could possibly play a role in ensuring that genes associated with H2AK119ub1 by PRC1 are also tri-methylated on H3K27 by PRC2 (Yuan et al., 2011).

In summary, we provide *in vitro* and *in vivo* evidence for the possible role of KDM2B as an H3K79me2/3 demethylase and as a co-repressor that regulates gene transcription through SIRT1-mediated chromatin silencing. Our model suggests that dynamic reversible regulation of histone methylation is indeed applied to H3K79 methylation. This model supports the existence of H3K79 demethylation by KDM2B and its role in transcriptional regulation (Figure 6E).

## Materials and methods

### DNA constructs

EFplink2-FLAG-KDM2B was acquired. The full-length KDM2B coding sequence (CDS) was transferred into p3XFLAG-CMV™-10, and a catalytic mutant construct was generated by H242A point mutation. The region (amino acids 1–734) containing the JmjC, CXXC zinc finger, and PHD domains was cloned into pGEX-4T1 for protein purification. pECE-FLAG-SIRT1 was purchased from Addgene (#1791). The SIRT1 CDS was subcloned into the modified pcDNA6-HA-MYC-HIS (Invitrogen).

### Cell culture

K562 cells were grown in Roswell Park Memorial Institute (RPMI) 1640 medium. 293T cells were grown in Dulbecco’s modified Eagle’s medium (DMEM) containing 10% heat-inactivated fetal bovine serum and 0.05% penicillin-streptomycin, at 37 °C in a 5% CO_2_ atmosphere. 50 μM sirtinol (Sigma-Aldrich) was treated for 2 h. 100 μM PRT4165 (Tocris Bioscience) was treated for 2 h in serum-free medium.

### Antibodies

Antibodies used in this study: anti-KDM2B (Millipore), anti-SIRT1 (Millipore), anti-DOT1L (Santa Cruz Biotechnology), anti-β actin (Santa Cruz Biotechnology), anti-FLAG (Sigma-Aldrich), anti-hemagglutinin (HA) (Santa Cruz Biotechnology), anti-H3K79me3 (Abcam or Epigentek), anti-H3K79me2 (Abcam), anti-H3K79me1 (Abcam), anti-H3K36me2 (Abcam), anti-H3K36me3 (Upstate), anti-H3K4me3 (Abcam), anti-H4K16ac (Millipore), and anti-histone H3 (Santa Cruz Biotechnology).

### Peptide pull-down assay

Biotinylated dimethyl H3K79 peptides, H-RLVREIAQDFK[me2]TDLRFQSSAVK[biotin]-OH, and unmodified H3K79 peptides, H-RLVREIAQDFKTDLRFQSSAVK[biotin]-OH, were purchased from AnaSpec (California, United States). 5 μg peptides were pre-bound to streptavidin-Sepharose beads (GE Healthcare) and incubated overnight at 4°C with nuclear extract from K562 cells, dialyzed against dialysis buffer (20 mM HEPES, pH 7.9, 1.5 mM MgCl_2_, 20% glycerol, 0.2 mM EDTA, 100 mM KCl, and 0.5 mM DTT), and pre-cleared. The supernatant was stored, and the particles were washed five times with 1 mL of washing buffer (20 mM HEPES, pH 7.9, 1.5 mM MgCl_2_, 20% glycerol, 0.2 mM EDTA, 100 mM KCl, 0.5 mM DTT, and 0.1% Triton X-100). Peptide-binding proteins were separated by SDS-PAGE. A portion of them was stained with silver, while the rest was divided into three parts to be analyzed by LC-MS/MS.

### In-gel protein digestion and liquid chromatography tandem-mass spectrometry (LC-MS/MS)

Mass spectrometry and proteomic analyses were carried out at the Korea Basic Science Institute. Separated proteins by two-dimensional gel electrophoreses (2-DE) were visualized using the PlusOne Silver Staining Kit (Amersham Biosciences), according to the manufacturer’s protocol. After electrical scanning and analysis of silver-stained gels using Phoretix Expression software ver. 2005 (Nonlinear Dynamics), the protein bands of interest were excised and digested in-gel with sequencing grade modified trypsin (Promega, Madison, WI, USA). Briefly, excised protein bands were washed with a 1:1 mixture of acetonitrile and 25 mM ammonium bicarbonate (pH 7.8), and subsequently dried using a SpeedVac concentrator. After drying, rehydration was performed with 25 mM ammonium bicarbonate (pH 7.8) and trypsin. Tryptic peptides were extracted from supernatants with a 50% aqueous acetonitrile solution containing 0.1% formic acid. After preparation, tryptic peptides were analyzed using reversed-phase capillary high-performance liquid chromatography (HPLC) directly coupled to a Finnigan LCQ ion trap mass spectrometer. Three extractions were performed to recover all tryptic peptides from the gel slices. Recovered peptides were concentrated by drying the combined extracts in a vacuum centrifuge. Concentrated peptides were mixed with 20 μL of 0.1% formic acid in 3% acetonitrile. Nano LC of the tryptic peptides was performed using the Waters Nano LC system, equipped with a Waters C18 nano column (75 μm × 15 cm, nanoAcquity UPLC column). Binary solvent A1 contained 0.1% formic acid in water, and binary solvent B1 contained 0.1% formic acid in acetonitrile. Samples (5 μL) were loaded onto the column, and peptides were subsequently eluted with a binary solvent B1 gradient (2–40%, 30 min, 0.4 μL/min). The lock mass, [Glu1] fibrinopeptide at 400 fmol/μL, was delivered from the auxiliary pump of the Nano LC system at 0.3 μL/min to the reference sprayer of the NanoLockSpray source.

### Stable knockdown cell lines

DNA oligonucleotides encoding KDM2B short hairpin RNA (shRNA) #1 (5′-CTGAACCACTGCAAGTCTATC-3′) and KDM2B shRNA #2 (5′-CGGCCTTTACAAGAAGACATT-3′) were subcloned into the pLKO.1-puro lentiviral vector (Addgene), according to standard procedures. To produce virus particles, 293T cells were co-transfected with plasmids encoding vesicular stomatitis virus glycoprotein (VSV-G), NL-BH, and the shRNAs. Two days after transfection, the supernatants containing the viruses were collected and used to infect 293T cells in the presence of polybrene (8 μg/mL). After lentiviral infection of the 293T cells, the addition of puromycin (1 mg/mL) selected for cells stably expressing shRNAs of KDM2B.

### Immunofluorescence analysis

293T cells were transfected with the p3XFLAG-CMV^TM^-KDM2B wild-type or the catalytic mutant, using Lipofectamine 2000 reagent (Invitrogen). The cells were fixed 48 h after transfection, permeabilized, and stained with the appropriate antibodies and secondary fluorochrome-labeled antibodies. The stained cells were analyzed by confocal laser fluorescence microscopy. The following antibodies were used: anti-FLAG, anti-H3K79me3 (Epigentek), anti-H3K79me1, goat anti-mouse IgG-h+l fluorescein isothiocyanate (FITC) conjugated (Bethyl Laboratories), Cy™3-conjugated AffiniPure goat anti-rabbit IgG (H+L) (Jackson Immunoresearch).

### Histone acid extraction and nucleosome extraction

To extract histones, cell pellets were resuspended in PBS with 0.5% Triton X-100 and protease inhibitors, and the tubes were subsequently incubated at 4°C for 30 min to lyse the cells. The lysates were centrifuged at 4°C for 10 min at 10,000 × *g*, and the pellets were resuspended in 0.4 N H_2_SO_4_. The samples were centrifuged at 4°C for 10 min at 16,000 x *g*. The pellets were again resuspended in 100% trichloroacetic acid (TCA) and centrifuged at 4°C for 10 min at 16,000 × *g*. The histone-containing pellets were collected and eluted in distilled water. To extract nucleosomes, cell pellets were resuspended in RSB buffer (10 mM Tris-HCl, pH 7.4, 10 mM NaCl, 3 mM MgCl_2_, 0.5% NP-40, 1 mM DTT, and 1 mM PMSF), and the tubes were subsequently incubated at 4°C for 30 min to lyse the cells. The lysates were centrifuged at 4°C for 5 min at 4,000 rpm, and the pellets were resuspended in RSB buffer. The samples were sonicated and centrifuged at 4°C for 5 min at 4,000 rpm. The pellets were washed four times with RSB buffer and eluted in the same buffer.

### *In vitro* histone demethylase assay

Bulk histones (Sigma-Aldrich), nucleosomes extracted from 293T cells, and methyl lysine analog (MLA) containing recombinant H3K79me3 histones (Active Motif) (Jia et al., 2009; Simon et al., 2007) were incubated with purified glutathione S-transferase (GST), GST-KDM2B_1–734_, or GST-KDM2A_1–687_, overnight at 37°C in demethylation assay buffer (20 mM Tris-HCl, pH 7.3, 150 mM NaCl, 1 mM alpha-ketoglutarate, 50 μM FeSO_4_, and 2 mM ascorbic acid).

### Formaldehyde dehydrogenase (FDH) assay

Formaldehyde formation was measured by a spectrophotometric assay (Lizcano et al., 2000) using FDH. GST-KDM2B and MLA histone H3K79me3 were first incubated for 3 hours at 37°C in demethylation assay buffer. The reaction solution was immediately mixed in buffer containing 50 mM potassium phosphate, pH 7.2, 2 mM NAD+, and 0.1 U FDH. The FDH reaction was initiated and absorbance at 340 nm was recorded for 5 minutes. The absorbance at 340 nm (∊340 = 6.22 mM–1cm–1 for NADH) was measured at each time point in a 0.5 min interval using WPA Lightwave S2000 UV/Vis spectrophotometer. The OD 340 nm absorbance at the moment of the FDH addition was considered as 0 and this was used as the 0 min time point. The data were analyzed using the Excel program.

### Nano LC-LTQ-Orbitrap Elite analysis

Digested plasma samples were dissolved in mobile phase A and analyzed using an LC-MS/MS system consisting of a Nano Acquity UPLC system (Waters, USA) and an LTQ Orbitrap Elite mass spectrometer (Thermo Scientific, USA) equipped with a Nano-electrospray source. An autosampler was used to load 5μL aliquots of the peptide solutions into a C18 trap column of internal diameter (ID) 180 μm, length 20 mm, and particle size 5 μm (Waters, USA). Peptides were desalted and concentrated on the trap column for 10 min, at a flow rate of 5 μL/min. The trapped peptides were then back-flushed and separated on a homemade microcapillary C18 column of ID 100 μm and length 200 mm (Aqua; particle size 3 μm, 125 Å). The mobile phases were composed of 100% water (A) and 100% acetonitrile (ACN) (B), each containing 0.1% formic acid. The LC gradient began with 5% mobile phase B and was maintained for 15 min. Mobile phase B was linearly ramped to 15% for 5 min, 50% for 75 min, and 95% for 1 min. Then, 95% mobile phase B was maintained for 13 min before decreasing to 5% for another 1 min. The column was re-equilibrated with 5% mobile phase B for 10 min before the next run. The voltage applied to produce the electrospray was 2.2 kV. During the chromatographic separation, the LTQ Orbitrap Elite was operated in data-dependent mode. MS data were acquired using the following parameters: full scans were acquired in the Orbitrap at a resolution of 120,000 for each sample; six data-dependent collision-induced dissociation (CID) MS/MS scans were acquired per full scan; CID scans were acquired in a linear trap quadrupole (LTQ) with 10 ms activation times performed for each sample; 35% normalized collision energy (NCE) was used in CID; and a 2 Da isolation window for MS/MS fragmentation was applied. Previously fragmented ions were excluded for 180 s. LC-MS/MS was performed using Nano LC-LTQ-Orbitrap Elite mass spectrometry at the Korea Basic Science Institute (Ochang Headquarters, Division of Bioconvergence Analysis).

### Isothermal titration calorimetry (ITC)

Isothermal titration calorimetry (ITC) experiments were carried out using an Auto-iTC200 Microcalorimeter at Korea Basic Science Institute. Tri-methyl H3K79 peptides, AQDFK(me3)TDLR, and unmodified H3K79 peptides, AQDFKTDLR, were purchased from AnyGen. Protein GST-KDM2B was purified and prepared in the sample cell, and the ligand (H3K79me3 peptide or H3K79me0 peptide) was loaded into the injectable syringe. All samples were prepared in 1X PBS. Titration measurements that consisted of 19 injections (2 μL) with 150 s spacing were performed at 25 °C, while the syringe was stirred at 700 rpm. The data were analyzed using the MicroCal Origin™ software.

### Radioactive assay

Histone methyltransferase (HMTase) assays were carried out at 30°C overnight, in 30μL volume containing 20 mM Tris-HCl [pH 8.0], 4 mM EDTA, 1 mM phenylmethylsulfonyl fluoride (PMSF), 0.5 mM dithiothreitol (DTT), 100 nCi of S-Adenosyl-L-methionine [methyl-14C] (^14^C-SAM) (Perkin Elmer), 15 μg core histones from calf thymus (Sigma-Aldrich), and GST-DOT1L, GST-MMSET, or GST-SMYD3, respectively. ^14^C-labeled histones were subjected to histone demethylase assay using either purified GST or GST-KDM2B_1–734_. The histones were transferred onto p81 filter paper (Upstate) and washed three times with 95% ethanol for 5 min at room temperature. The filters were allowed to air dry, and 2 mL of Ultima Gold (Perkin Elmer) was added afterwards. ^14^C-SAM was then quantified using a scintillation counter.

### Luciferase assay

Luciferase assays were conducted using the *HOXA7*, *MEIS1*, simian virus 40 (SV40), and thymidine kinase (TK) promoter reporter systems. 293T cells were co-transfected with the corresponding promoter reporter constructs and the indicated DNA constructs, using polyethylenimine (PEI). Cells were harvested after 48 h and assayed for luciferase activity using a luciferase assay system (Promega). Each value is expressed as the mean of five replicates of a single assay. All experiments were performed at least three times.

### Chromatin immunoprecipitation (ChIP)-quantitative polymerase chain reaction (qPCR) and ChIP sequencing (ChIP-Seq) Analyses

Formaldehyde (1%) was added to the medium for 10 min at room temperature, followed by the addition of 125 mM glycine for 5 min at room temperature. Adherent cells were scraped from dishes into 1 mL PBS. The scraped cells were centrifuged, and the resulting pellets were washed once with PBS. The pellets were resuspended in sodium dodecyl sulfate (SDS) lysis buffer (1% SDS, 10 mM EDTA, and 50 mM Tris-HCl, pH 8.1). The cell lysates were sonicated, diluted with five volumes of dilution buffer (0.01% SDS, 1.2 mM EDTA, 1.1% Triton X-100, 167 mM NaCl, and 16.7 mM Tris-HCl, pH 8.1), and incubated overnight with indicated antibodies (1 μg antibody for each IP reaction). The next day, protein A/G-Agarose beads (GenDEPOT) were added to the reaction, incubated for 2 h, and washed with low salt wash buffer, high salt wash buffer, LiCl immune wash buffer, and Tris-EDTA (TE) buffer. The immunoprecipitates were eluted and reverse cross-linked at 65°C. Afterwards, the DNA fragments were purified either for polymerase chain reaction (PCR) amplification or for sequencing using Illumina HiSeq 2000.

Values represent mean ±SD of technical duplicates from a representative experiment. All experiments were performed three times with similar results.

### qPCR Analyses

The immunoprecipitated DNA fragments were purified and PCR-amplified for quantification, using each PCR primer pair. Disassociation curves were generated after each PCR run to ensure amplification of a single product of the appropriate length. The mean threshold cycle (C_T_) and standard error values were calculated from individual C_T_ values, obtained from duplicate reactions per stage. The normalized mean C_T_ value was estimated as ΔC_T_ by subtracting the mean C_T_ of the input. To analyze promoter regions of *HOXA7*, *MEIS1*, *ALX1*, and *GATA4*, and 5’ ends of *PDE3B*, primer sets indicated in Table S1 were used. The primer concentration used for qPCR (Bio-Rad) was 0.2 μM/10 μL. The thermal cycler conditions were as follows: 15 min of holding at 95°C, followed by 39 cycles at 94°C for 15 s, 56°C for 30 s, and 72°C for 30 s.

### Statistics

Data were analyzed by Student t-test as indicated.

## Acknowledgements

We are grateful to Dr. Vivian Bardwell (University of Minnesota) for providing EFplink2-FLAG-KDM2B. We would like to thank Dr. Jin Young Kim and Dr. Ju Yeon Lee (Korea Basic Science Institute, Ochang Headquarter, Division of Bioconvergence Analysis) for the nano LC-LTQ-Orbitrap analysis. The ChIP-seq data are available from GEO (Gene Expression Omnibus) under accession number GSE89052.

## Additional information

National Research Foundation of Korea (NRF) grant funded by the Ministry of Science and Technology, Basic Science Research program [NRF-2017R1A2B4004407 to J.Y.K. and J.Y.H.]; National Research Foundation of Korea (NRF) grant funded by the Ministry of Science, ICT & Future Planning [NRF-2016R1A4A1008035 to J.Y.K., J.W.P., H.C., J.Y.H., and Y.C.C.]; This research was supported by the Chung-Ang University *Excellent Student Scholarship* in 2013.

